# Lymphatic Dysfunction Models an Autoimmune Emphysema Phenotype of Chronic Obstructive Pulmonary Disease

**DOI:** 10.1101/2023.10.31.564938

**Authors:** Barbara Summers, Kihwan Kim, Tyler M. Lu, Sean Houghton, Anjali Trivedi, Joselyn Rojas Quintero, Juan Cala-Garcia, Tania Pannellini, Francesca Polverino, Raphaël Lis, Hasina Outtz Reed

## Abstract

Chronic Obstructive Pulmonary Disease (COPD) is a heterogeneous disease that is characterized by many clinical phenotypes. One such phenotype of COPD is defined by emphysema, pathogenic lung tertiary lymphoid organs (TLOs), and autoantibody production. We have previously shown that lymphatic dysfunction can cause lung TLO formation and lung injury in mice. We now sought to uncover whether underlying lymphatic dysfunction may be a driver of lung injury in cigarette smoke (CS)-induced COPD. We found that lung TLOs in mice with lymphatic dysfunction produce autoantibodies and are associated with a lymphatic endothelial cell subtype that expresses antigen presentation genes. Mice with underlying lymphatic dysfunction develop increased emphysema after CS exposure, with increased size and activation of TLOs. CS further increased autoantibody production in mice with lymphatic dysfunction. B-cell blockade prevented TLO formation and decreased lung injury after CS in mice with lymphatic dysfunction. Using tissue from human COPD patients, we also found evidence of a lymphatic gene signature that was specific to patients with emphysema and prominent TLOs compared to COPD patients without emphysema. Taken together, these data suggest that lymphatic dysfunction may underlie lung injury in a subset of COPD patients with an autoimmune emphysema phenotype.

## Introduction

Chronic obstructive pulmonary disease (COPD) is a clinically and pathologically heterogeneous disease. Though commonly caused by cigarette smoke (CS), the mechanisms that lead some patients and not others to have progressive to disease are not entirely clear. Furthermore, we are just beginning to understand the biologic endotypes that drive distinct clinical phenotypes of this disease. Given that there are currently no disease-modifying therapies for COPD, understanding the distinct molecular pathways that drive lung injury in COPD promises to lead to novel therapeutic strategies that are specifically targeted to patients’ underlying pathobiology.

Tertiary lymphoid organs (TLOs) are a prominent histologic finding in a clinical phenotype of COPD that is characterized by emphysema and an adaptive immune signature that promotes autoantibodies and autoreactive T cells (1–3). Though an ‘autoimmune emphysema’ phenotype of COPD is present in a subset of patients, it is unclear what are the underlying drivers of their specific disease pathogenesis. However, the pathogenic nature of the TLOs that are prominent in autoimmune emphysema have been the subject of a great deal of investigation (2, 4–6). TLOs resemble lymph nodes in their structure and organization and play protective or pathogenic roles in lung disease that are context dependent. In CS-induced emphysema, TLOs are thought to be pathogenic due at least in part to the generation of self-reactive immune responses to antigens that are generated as a result of CS-mediated lung damage (2). The lymphatic vasculature mediates both leukocyte trafficking and antigen drainage in the lung and are found near TLOs when they form (7). We have previously found that lymphatic dysfunction alone is sufficient to induce formation of lung TLOs in mice (8), however, whether the lymphatics play a role in the pathogenic function of TLOs in the setting of CS-induced emphysema is unknown.

We sought to uncover whether lymphatic dysfunction drives lung injury after CS exposure using a mouse model of COPD. We found that lung TLOs in mice with lymphatic dysfunction are associated with autoantibody production and a lymphatic endothelial cell (LEC) subtype characterized by expression of antigen presentation genes. Mice with lymphatic dysfunction developed increased emphysema after CS exposure, which was associated with increased activation of TLOs and further autoantibody production. B-cell blockade in mice with lymphatic dysfunction prevented TLO formation and decreased lung injury after CS exposure. Interestingly, we also found that expression of lymphatic endothelial cell markers is seen in lung tissue from COPD patients with an emphysema phenotype characterized by prominent TLOs, and not in COPD patients without emphysema. Taken together, these data suggest that lymphatic dysfunction may play a role in an autoimmune emphysema phenotype of COPD.

## Results

TLOs in mice with lymphatic dysfunction are associated with autoantibody production C-type lectin-like type II (CLEC2) is a platelet receptor that is necessary for lymphatic function due to its role during development in maintaining separation between the venous and lymphatic vasculature. In the absence of CLEC2, there is primary formation of lymphatic vessels but defects in remodeling and vessel maturation, with impaired lymph flow due to retrograde flow of blood into the lymphatic system that results in edema, chylothorax, lack of lung inflation, and other sequelae of impaired lymphatic function (8–11). We have shown that CLEC2-deficient mice that survive to adulthood have systemically impaired lymph flow, which in the lung results in impaired leukocyte trafficking, TLO formation, and a lung injury phenotype that resembles human emphysema (8). To better understand the role of TLOs formed due to lymphatic dysfunction on the development of emphysema, we generated mice with platelet-specific loss of CLEC2 (*Clec2^fl/fl;^PF4Cre*, hereafter *Clec2^pltKO^*) that have less severe lymphatic dysfunction than global loss of CLEC2, likely due to incomplete recombination in megakaryocytes (9). *Clec2^pltKO^* mice are born at expected frequencies and survive to adulthood, but still have evidence of lymphatic impairment and develop lung TLOs (Figure 1a,b). These TLOs consist predominantly of B cells, including plasma cells (Figure 1c-f), and are not typically seen in the lungs of control mice in the absence of inflammation (6, 8). TLOs in *Clec2^pltKO^* mice were associated with an increase in total IgA and IgG antibodies in the bronchoalveolar lavage (BAL) fluid of these animals (Figure 1g,h), consistent with the increased presence of B cells in the lungs. Because autoantibodies to extracellular matrix (ECM) proteins are seen in a subset of COPD patients with autoimmune emphysema that have prominent TLOs (12–14), we assessed whether these autoantibodies were also present in *Clec2^pltKO^* mice. Interestingly, we found that autoantibodies to collagen and elastin were significantly increased in the BAL fluid of *Clec2^pltKO^* mice compared to control (Figure 1i-k). This appeared to be a result of lung TLOs rather than systemic dysfunction, as we did not see increased anti-elastin in the serum of these mice (Figure 1l). To determine whether these autoantibodies are also seen in other settings of spontaneous lung TLO formation due to lymphatic dysfunction, we analyzed the lungs of CCR7-deficient mice, which have a lymphatic trafficking defect and develop lung TLOs due to defective dendritic cell migration mediated by CCL21 on the lymphatic endothelium (15). We found that CCR7-deficient mice develop lung TLOs and alveolar enlargement resembling emphysema, similarly to *Clec2^pltKO^*mice (Supplemental Figure 1a-c), which has been reported in previous studies (16). We also detected autoantibodies in the lungs of these mice (Supplemental Figure 1d-f), supporting previous reports of an autoimmune phenotype in these animals (17), suggesting that TLOs that form due to lymphatic dysfunction may result in autoimmunity despite differing mechanisms of lymphatic impairment.

**Figure 1:**
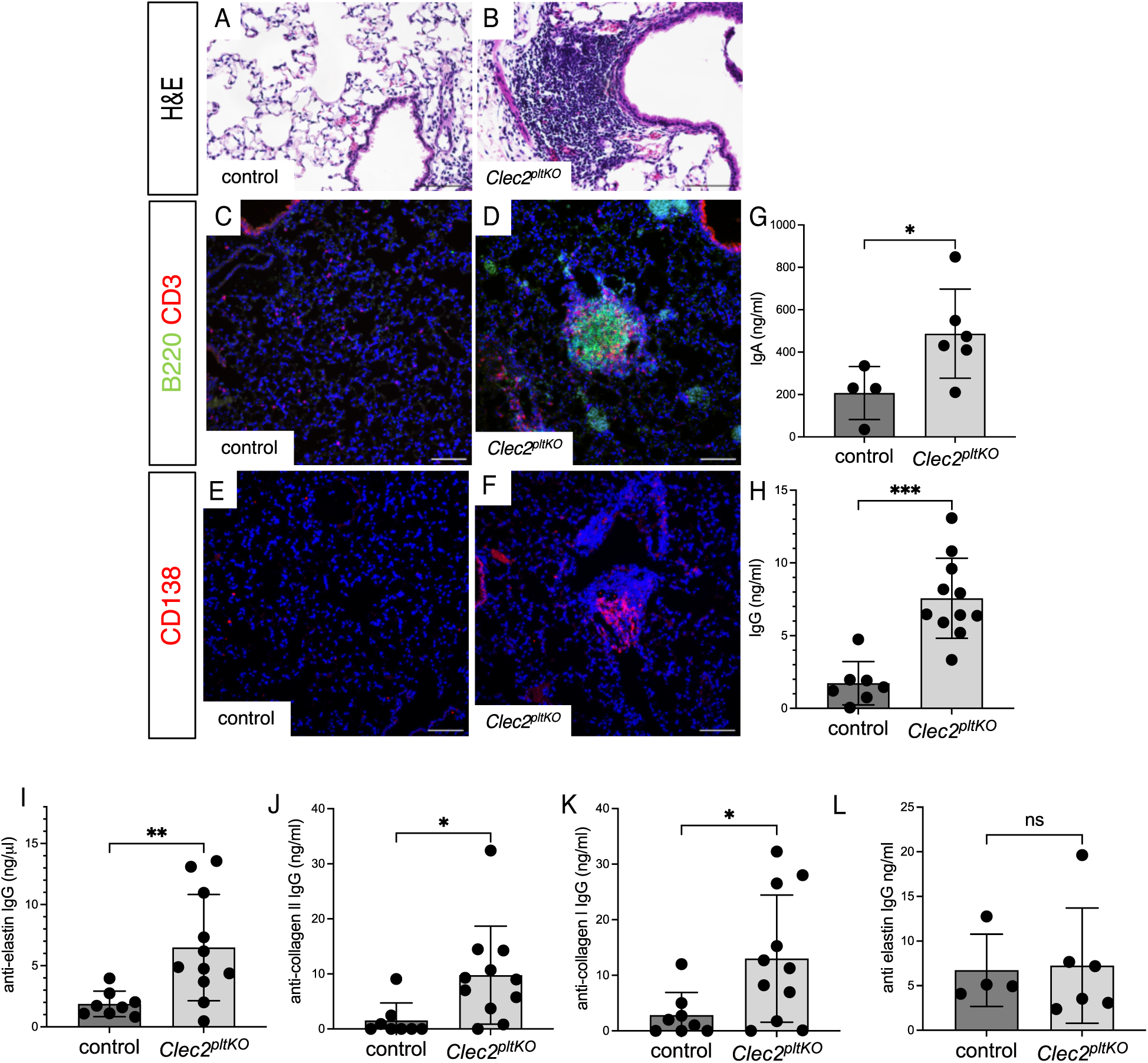
**TLOs in mice with lymphatic dysfunction are associated with autoantibodies to extracellular matrix proteins** A, B: H&E staining of lung tissue from control (A) and *Clec2^pltKO^* (B) mice. C-F: Immunohistochemical staining of lung tissue from control (C,E) and *Clec2^pltKO^* (D,F) mice. Slides were stained with antibodies for B220 and CD3 (C,D) or CD138 (E,F) and counterstained with DAPI. G, H: ELISA for total IgA and IgG in BAL fluid from control and *Clec2^pltKO^* mice. I-K: ELISA for anti-elastin IgG (I), anti-collagen II IgG (J), and anti-collagen I IgG (K) in BAL fluid from *Clec2^pltKO^* mice. L: ELISA for anti-elastin IgG in mouse serum. All images are representative of at least 8 mice from 2 separate experiments. Scale bars = 100 µm. All values are means ± SEM. *P* value calculated by Student’s *t* test. *\*P* < 0.05, \*\**P*< 0.01, \*\*\**P*< 0.001.

Single cell RNA Sequencing Reveals Unique LEC Subtype in Mice with TLOs due to Lymphatic Dysfunction Lymphatic vessels are closely associated with TLOs, and changes in lymphatic function or lymphatic leukocyte trafficking results in TLO formation in the lung and elsewhere in multiple mouse models (8, 15, 18, 19). We sought to determine whether the lung lymphatic endothelium is altered in mice with lymphatic dysfunction that form TLOs. We performed single cell RNA sequencing on isolated lung LECs from control and *Clec2^pltKO^* mice and identified 6 distinct cell cultures of lung LECs across all samples (Figure 2a,b). We found a unique LEC subtype in that was unique to *Clec2^pltKO^* mice (cluster 2) that was characterized by increased expression of genes important for antigen presentation, including the MHC II invariant chain CD74 and class II histocompatibility genes H2-Aa1, H2-Ab1, and H2-Eb1 (Figure 2c,d). Pathway analysis also demonstrated enrichment of MHC II protein complex binding in cluster 2 that is overwhelmingly unique to lung LECs from *Clec2^pltKO^* mice (Figure 2e). Flow cytometry confirmed surface expression of MHC II on a subset of LECs from the lungs of *Clec2^pltKO^* mice and not in control mice (Figure 2f). This immunomodulatory subtype lacks costimulatory molecules that are typically found in antigen presenting cells (20) and has also been seen in other settings of impaired lymphatic function where TLOs form (18), suggesting that the presence of this LEC subtype may be a marker of impaired lymphatic drainage and inflammation in multiple settings.

**Figure 2:**
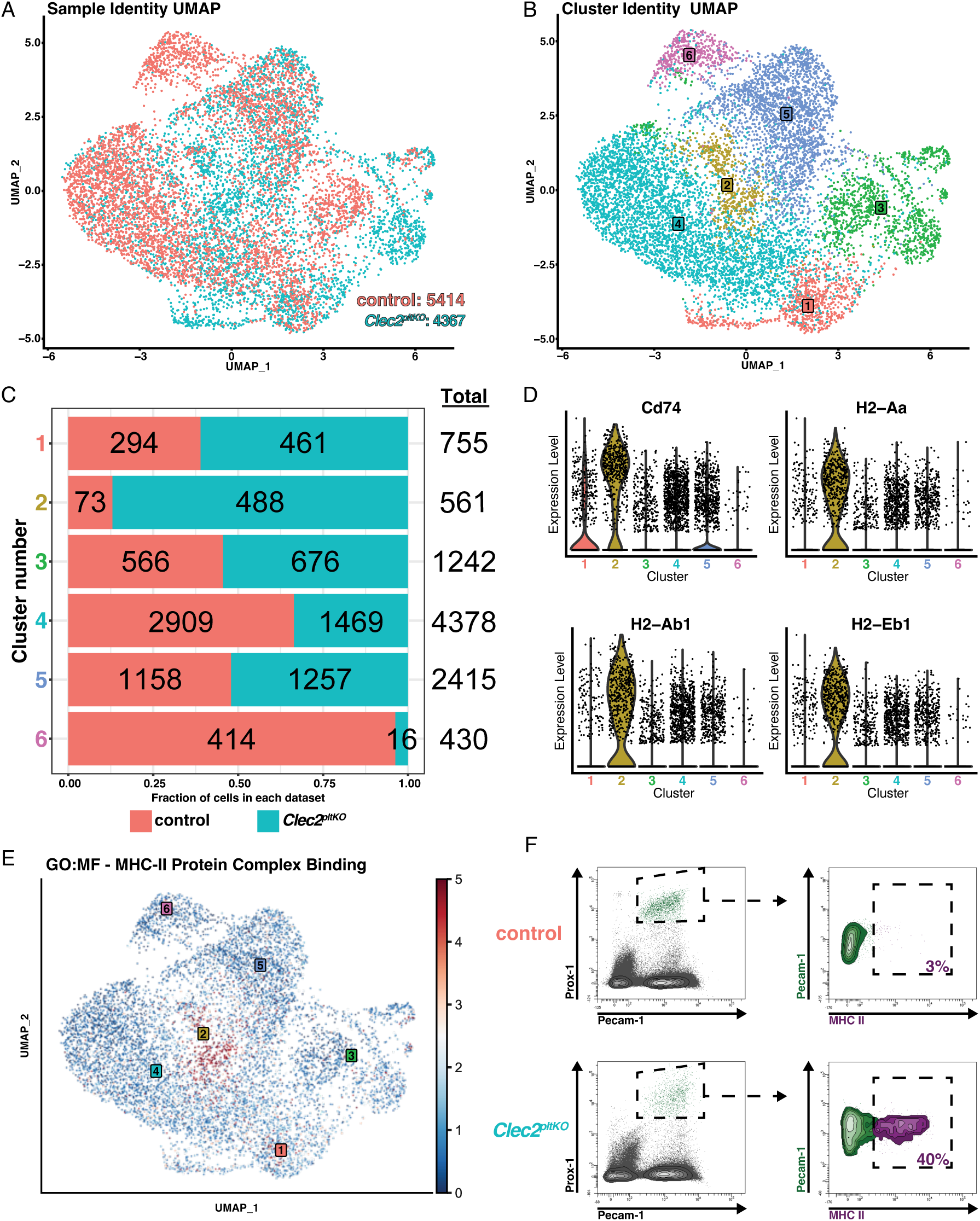
Single cell RNA sequencing reveals unique LEC subtype in mice with lymphatic dysfunction. A: Sample identity UMAP plot of control and *Clec2^pltKO^* mouse lung lymphatic endothelial cell (LEC) samples from n=3 biological replicates per group totaling 5414 control cells and 4367 *Clec2^pltKO^* cells. B: Cluster identity UMAP plot demonstrating 6 distinct cell clusters across all the LECs in our dataset. C: Cluster composition plot showing the distribution of control and *Clec2^pltKO^* LECs in each distinct cluster as well as the total cell number per cluster which highlights cluster 2 as a predominantly *Clec2^pltKO^* cluster. D: Violin plots demonstrating significant di erential expression of select MCH II pathway genes with cluster 2 showing the highest expression across all markers. E: Pathway enrichment UMAP plot demonstrating the expression of MHC II Protein Complex Binding (GO:0023026) Molecular Function gene signatures across all cells revealing enrichment in the *Clec2^pltKO^*dominant cluster 2. F: Flow cytometry plots showing PROX1^+^ Pecam-1^+^ LECs (Ptprc^-^ Lin^-^) pooled from control (n = 5) vs *Clec2^pltKO^* (n = 3) mice.

### Mice with Lymphatic Dysfunction Develop Exacerbated Emphysema after CS Exposure

We next tested whether lymphatic dysfunction in *Clec2^pltKO^* mice changes the response to CS in a model of COPD. While control mice develop only mild emphysema after chronic CS exposure in this model (21, 22), *Clec2^pltKO^*mice developed increased emphysema as evidenced by histology and quantification of alveolar enlargement (Figure 2a-e). This was associated with alveolar injury, with elastin breakdown seen in lung lysates from control *Clec2^pltKO^* mice and those exposed to CS (Figure 3f). CS-exposed *Clec2^pltKO^* mice also had an increase in both the size and number of lung TLOs compared to both RA *Clec2^pltKO^* mice as well as control mice exposed to CS (Figure 3g-p). This correlated with an increase in CXCL13, CCL5, CXCL9, and IL-16 (Supplemental Figure 2), cytokines that have previously been shown to be associated with recruitment of leukocytes, formation and maintenance of TLOs, and B-cell/T-cell crosstalk (23–27). *Clec2^pltKO^*mice have increased lymphatic vessel staining in the lung parenchyma compared to control mice likely reflecting the abnormal morphology of these vessels (8), though this was not further increased by CS exposure (Figure 3q).

**Figure 3:**
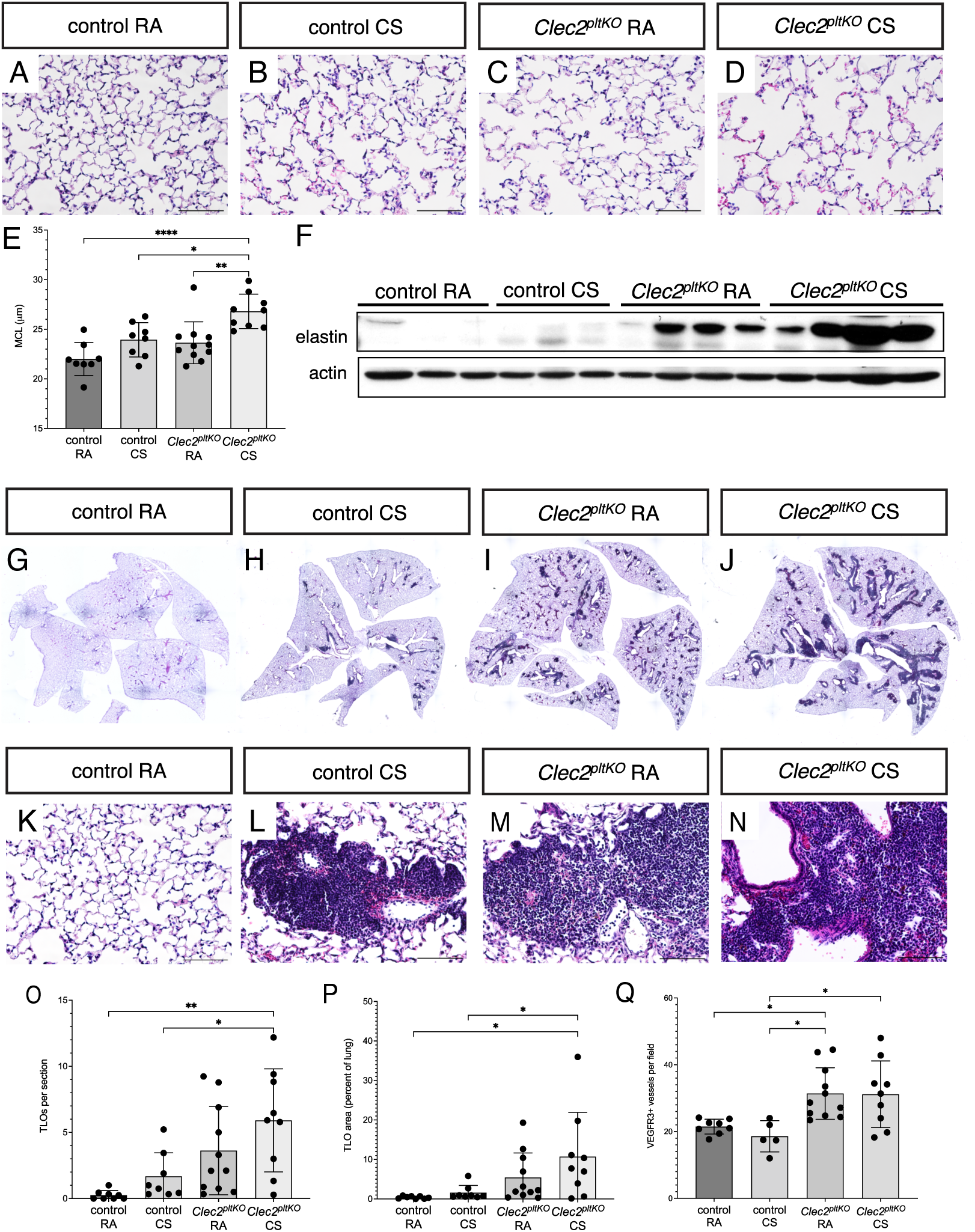
**Exacerbated Emphysema after CS exposure in mice with lymphatic dysfunction**A-D: H&E staining of lung sections from control and *Clec2^pltKO^* mice after 8 months of exposure to cigarette smoke (CS) or age-matched room air (RA) controls. E: Quantification of alveolar size after 8 months of CS exposure or RA by mean chord length (MCL). F: Western blot of lung lysates from mice exposed to 8 months of CS or RA for soluble elastin. Beta-actin (BA) was probed on the same membrane as a loading control. G-J: Whole lung images of H&E-stained sections from mice exposed to 8 months CS or RA. K-N: H&E images of TLOs in sections from mice exposed to CS or RA. O: Quantification of TLOs per whole lung section in control and *Clec2^pltKO^* mice exposed to 8 months CS or RA. P: Quantification of TLO area as a percentage of total lung area per section in control and *Clec2^pltKO^* mice exposed to 8 months CS or RA. Q: Quantification of lymphatic vessels, as measured by number of VEGFR3-positive vessels seen per field by immunohistochemistry. Scale bars = 100 µm. All values are means ± SEM. *P* value calculated by ANOVA. *\*P* < 0.05, \*\**P*< 0.01, \*\*\**P*< 0.001, \*\*\*\**P*< 0.0001.

### CS Promotes TLO Maturation

Given the increase in TLOs seen on histology with CS exposure, we next sought to determine whether CS altered TLO formation or composition. To quantitatively assess TLO composition and activation after CS exposure, we used GeoMx digital spatial proteomic profiling (NanoSpring) to compare TLOs formed from CS exposure in control *Clec2^pltKO^* mice, compared to TLOs in *Clec2^pltKO^* mice exposed to room air. We defined our regions of interest (ROIs) as the lung TLOs and used an antibody panel consisting of 28 targets toward both immunophenotyping as well as immune activation (Table 1). Interestingly, we detected only small differences in B cell, T cell, and myeloid markers in TLOs from these animals with the exception of significantly decreased CD8^+^ T cells in TLOs CS-exposed control mice compared to *Clec2^pltKO^* mice (Figure 4a-d). However, we found significantly increased MHC II presentation and CD40 in TLOs in both control and *Clec2^pltKO^* mice that were exposed to CS compared to TLOs in room air exposed *Clec2^pltKO^* mice (Figure 4a,b,e,f). Despite the difference in their number and size, TLOs in CS-exposed control and *Clec2^pltKO^* mice showed few statistically significant differences (Figure 4c), suggesting that CS exposure plays an important role in modulating TLO organization and activation even among TLOs that formed prior to CS exposure. MHC II presentation and CD40 engagement are required for germinal center formation and progression, as well as antibody isotype switching and generation of memory B cells (28). These results suggest that CS leads to activation and maturation of lung TLOs whether they are formed de novo or are preexisting as we result of lymphatic dysfunction.

**Figure 4:**
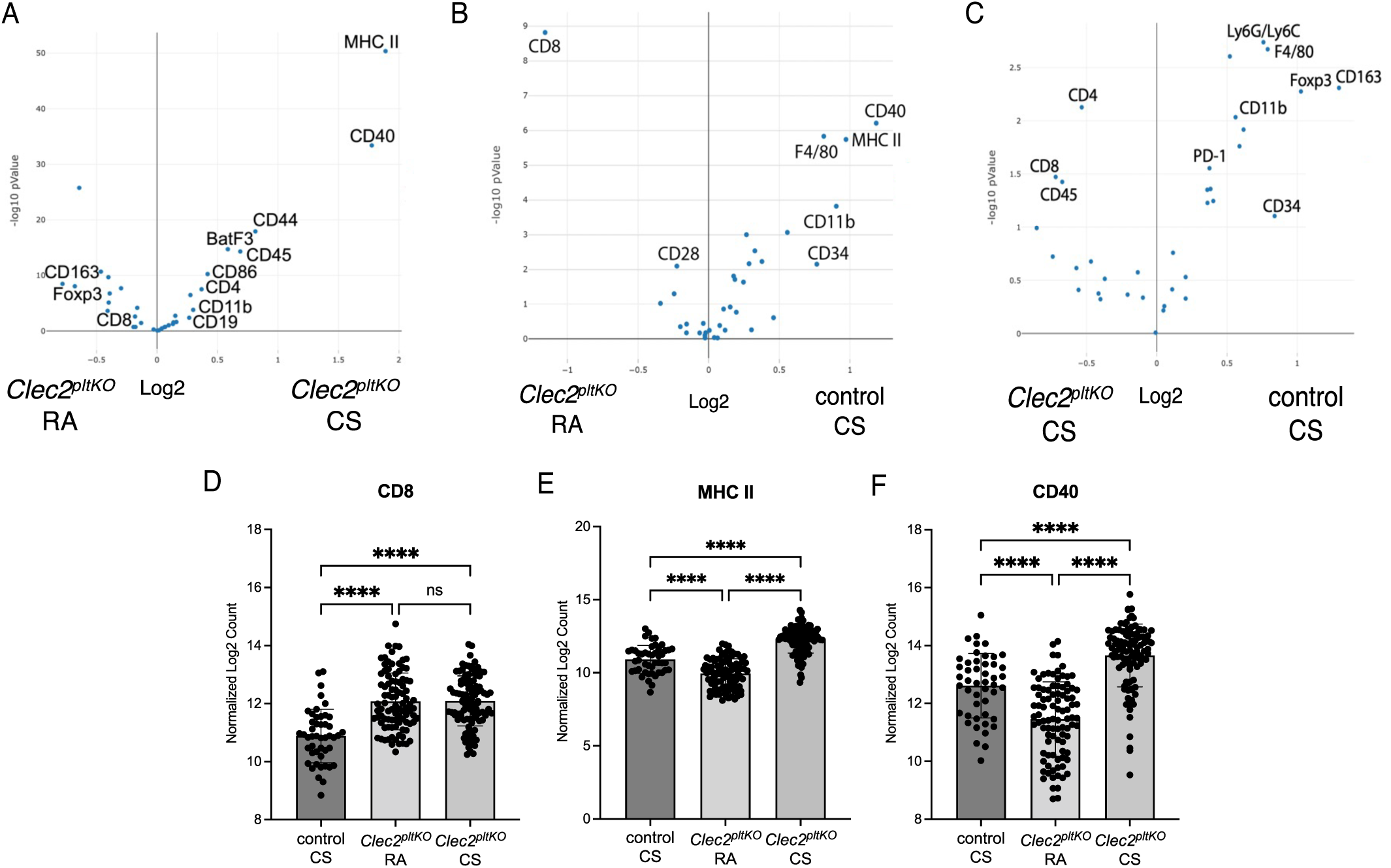
Increased TLO activation after CS exposure. A-C: Volcano plots showing protein expression of immune cell and activation markers TLOs from *Clec2^pltKO^* mice exposed to RA and CS, as well as control mice exposed to CS. Data were generated by NanoString GeoMX digital spatial profiling of 8-16 TLOs from n = 6 control CS, n= 7 *Clec2^pltKO^* RA mice, n = 7 *Clec2^pltKO^* CS mice. Log2 fold change is on x-axis, -log10 p-value on y axis. D-F: Quantification of CD8 (D), MHC II (E), and CD40 (F), in TLOs from *Clec2^pltKO^* mice exposed to RA and CS, as well as control mice exposed to CS by GeoMX digital spatial profiling. Each dot represents a ROI corresponding to a distinct TLO. \*\*\*\**P*< 0.0001, ns = not significant.

**Table 1:** Protein Targets for Spa4al Proteomic Analysis.

| Protein Name | Gene | Accession | Full Target Name | Protein ID |
| --- | --- | --- | --- | --- |
| CD127/IL7RA | Il7r | Q8C9W4,P16872,Q5FW78 | interleukin 7 receptor | DPROT_00036.1 |
| CD27 | Cd27 | B7ZW87,P41272,Q3U4X0,D3Z7W5 | CD27 antigen | DPROT_00089.1 |
| CD40L | Cd40lg | P27548,Q0VEI3 | CD40 ligand | DPROT_00088.1 |
| CD44 | Cd44 | A2APM2,A2APM4,A2APM3,P15379,A2APM1,Q80X37,Q3U8S1 | CD44 antigen | DPROT_00261.1 |
| CD86 | Cd86 | Q61238,P42082,Q549Q9 | CD86 antigen | DPROT_00273.1 |
| ICOS | Icos | Q5SUZ7,Q9WVS0,Q3V3X2 | inducible T cell co-stimulator | DPROT_00086.1 |
| PD-1 | Pdcd1 | Q02242,Q544F3 | programmed cell death 1 | DPROT_00065.1 |
| PD-L1 | Cd274 | Q3U472,Q9EP73 | CD274 antigen | DPROT_00080.1 |
| BatF3 | Batf3 | Q9D275 | basic leucine zipper transcription factor, ATF-like 3 | DPROT_00351.1 |
| CD19 | Cd19 | Q3TQG5,P25918 | CD19 antigen | DPROT_00073.1 |
| CD28 | Cd28 | P31041,Q8CDB3 | CD28 antigen | DPROT_00260.1 |
| CD34 | Cd34 | Q543B6,Q543Y2,Q64314 | CD34 antigen | DPROT_00048.1 |
| CD3e | Cd3e | A6H6M1,P22646 | CD3 antigen, epsilon polypeptide | DPROT_00074.1 |
| CD4 | Cd4 | Q3TSV7,P06332 | CD4 antigen | DPROT_00075.1 |
| CD8a | Cd8a | P01731,Q8CAX3 | CD8 antigen, alpha chain | DPROT_00078.1 |
| Fibronectin | Fn1 | A0A087WR50,Q3UHR1,Q3UGY5,Q3UHL6,P11276,A0A087WSN6,B9EHT6,A0A087WS56,Q3TBB4,Q3UZF9,B7ZNJ1,Q3UH17 | fibronectin 1 | DPROT_00081.1 |
| FOXP3 | Foxp3 | Q53Z59,Q99JB6 | forkhead box P3 | DPROT_00090.1 |
| GZMB | Gzmb | P04187,Q3TZH4 | granzyme B | DPROT_00079.1 |
| CD11b | Itgam | E9Q604,G5E8F1 | integrin alpha M | DPROT_00095.1 |
| CD11c | Itgax | Q9QXH4 | integrin alpha X | DPROT_00072.1 |
| CD14 | Cd14 | Q3UB18,Q4FJP7,P10810,Q3UE54 | CD14 antigen | DPROT_00091.1 |
| CD163 | Cd163 | Q2VLH6,B7ZMW6 | CD163 antigen | DPROT_00052.1 |
| CD39 | Entpd1 | Q8CDV7,Q544U5,P55772 | ectonucleoside triphosphate diphosphohydrolase 1 | DPROT_00105.1 |
| CD40 | Cd40 | P27512 | CD40 antigen | DPROT_00087.1 |
| CD68 | Cd68 | P31996 | CD68 antigen | DPROT_00436.1 |
| F4/80 | Adgre1 | Q61549,Q3U9R0 | adhesion G protein-coupled receptor E1 | DPROT_00071.1 |
| Ly6G/Ly6C | Ly6g,Ly6c1 | P0CW02,A0A087WRG9,A0A087WNZ5,A0A087WQF5 | lymphocyte antigen 6 complex, locus C1,lymphocyte antigen 6 complex, locus G | DPROT_00085.1 |
| MHC II | H2-Eb1,H2-Ab1 | Q3TD53,O78196,P14483,Q8BPA0 | histocompatibility 2, class II antigen A, beta 1,histocompatibility 2, class II antigen E beta | DPROT_00067.1 |

### CS Promotes Increased Autoantibody Production in Mice with Lymphatic Dysfunction

Autoantibody production is exacerbated by CS (29), and has been implicated in disease progression for patients with autoimmune emphysema (1, 3). Interestingly, these autoantibodies can be to a broad array of antigens, including ones that are not specific to the lung, and those that are more commonly seen in autoimmune diseases such as systemic lupus erythematosus SLE) and rheumatoid arthritis (RA) (30). We quantified autoantibody production after CS exposure in this model using an autoantigen microarray to detect autoantibodies to a broad range of proteins. We found that while IgG and IgA autoantibody production was broadly increased in the bronchoalveolar lavage (BAL) fluid of control room air exposed *Clec2^pltKO^*mice (Supplemental Figure 3), chronic CS exposure dramatically increased the presence of IgG autoantibodies within the lung tissue of *Clec2^pltKO^* mice (Figure 5a). CS-exposed *Clec2^pltKO^*mice had increased autoantibodies to a wide variety of antigens, including autoantibodies that are typically seen in autoimmune diseases (Figure 5b-g), such as Sm, SSA-A, Jo-1, Scl-70, and autoantibodies to complement proteins (3, 13, 14). We similarly found an increase in IgA autoantibodies in the lung tissue of CS-exposed *Clec2^pltKO^* mice (Supplemental Figure 4). To confirm that autoantibodies were preferentially localized to the lung tissue rather than in BAL after CS exposure, we detected widespread deposition of IgG antibodies throughout the lung tissue of CS-exposed *Clec2^pltKO^*compared to RA *Clec2^pltKO^*and control mice by immunohistochemistry (Figure 5h-k). These data suggest that *Clec2^pltKO^* mice have autoantibody production at baseline that is increased by CS exposure and preferentially localized to the lung tissue, where it would be predicted to be the most pathogenic.

**Figure 5:**
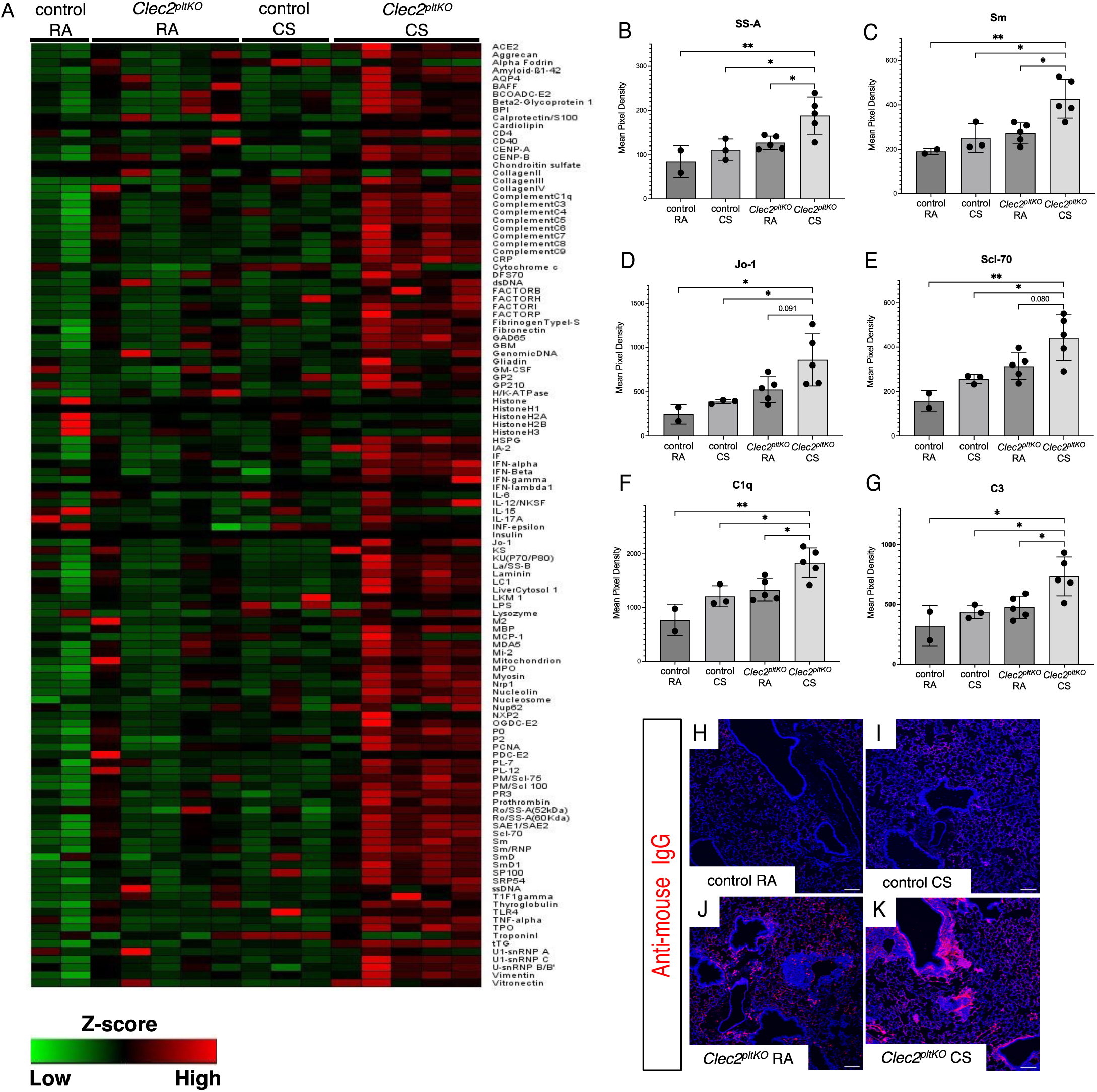
Increased autoantibodies after CS exposure in mice with lymphatic dysfunction. A: Heatmap of IgG autoantibodies to 128 antigens detected by ELISA using lung lysates from *Clec2^pltKO^* and control mice exposed to 8 months CS or RA and fluorescently labelled mouse anti-IgG. Fluorescence intensity was normalized to internal controls. B-G: Quantification of IgG autoantibodies commonly found in autoimmune disease including SS-A (A), Sm (B), Jo-1 (C), Scl-70 (E), and complement proteins C1q (F) and C3 (G). H-K: Detection of IgG deposition in lung tissue from *Clec2^pltKO^* and control mice exposed to 8 months CS or RA using fluorescently- labelled anti-mouse IgG. Images are representative of at least 6 mice. All values are means ± SEM. *P* value calculated by ANOVA. *\*P* < 0.05, \*\**P*< 0.01, \*\*\**P*< 0.001. Scale bars = 100 µm.

### B-Cell Blockade Prevents TLO Formation and Lung Injury after CS exposure in Mice with Lymphatic Dysfunction

We next addressed a causative role for TLOs formed due to lymphatic dysfunction in lung injury after CS exposure. We depleted B-cells in *Clec2^KO^* mice, who have global deletion of *Clec2* and severe lymphatic dysfunction leading to TLOs and lung injury (8), starting at 4 weeks of age using purified anti-mouse CD20 antibody. These mice were then subjected to chronic CS exposure, with ongoing B-cell depletion or injections with an isotype control antibody every 4 weeks (Figure 6a). We found that B cell blockade resulted in near complete inhibition of TLO formation in these animals (Figure 6b-g). As expected, B-cell blockade was also associated with markedly decreased antibody production in the lungs compared to mice treated with isotype control antibody (Supplemental Figure 5). Remarkably, inhibition of TLOs was associated with decreased alveolar enlargement (Figure 6H-J) and elastin breakdown (Figure 6K) after chronic CS exposure, despite continued lymphatic dysfunction in these animals. These data suggest that TLOs formed due to lymphatic dysfunction play a mechanistic role in lung injury after CS exposure.

**Figure 6:**
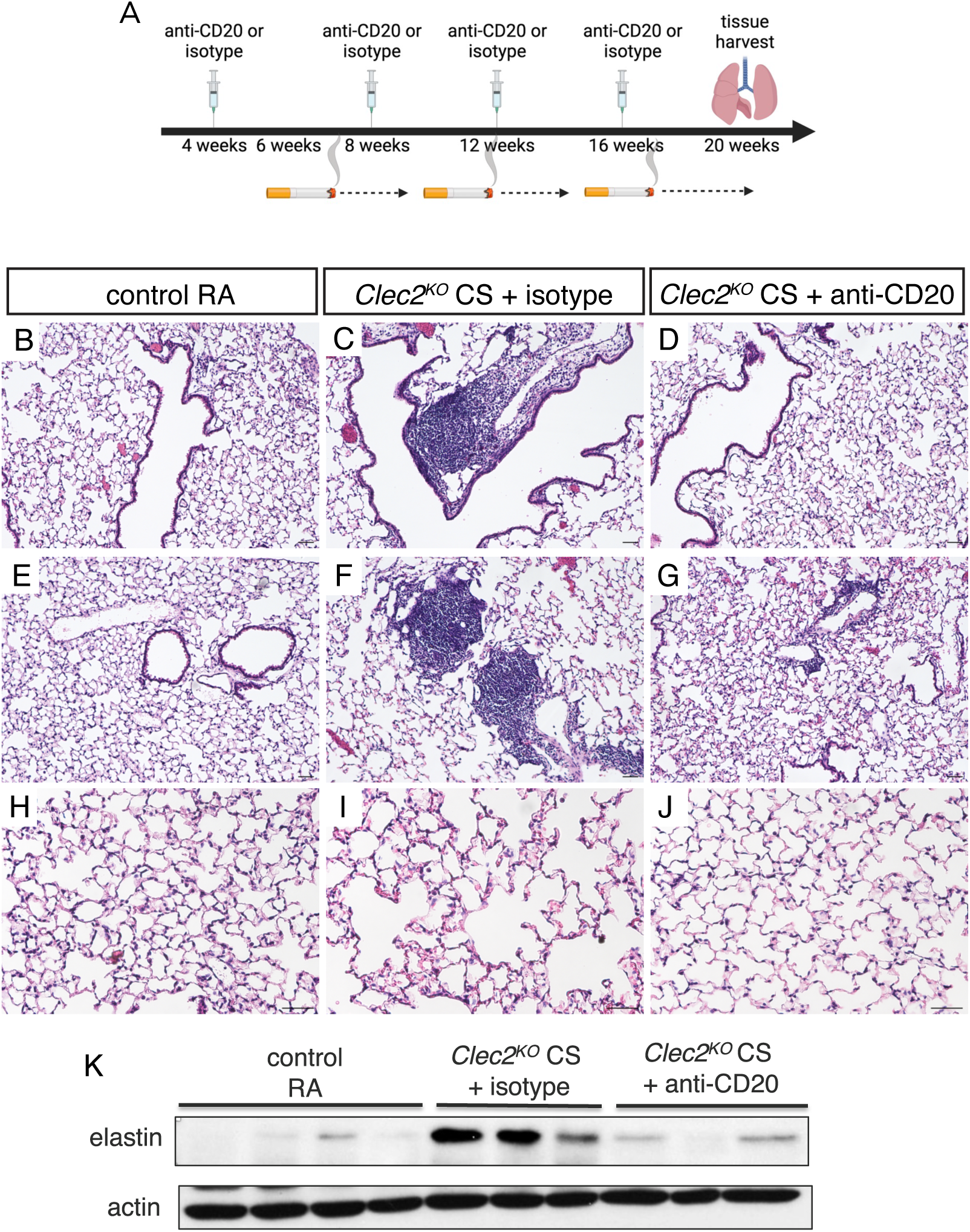
B-cell Blockade Prevents TLO Formation and Alveolar Enlargement in Mice with Lymphatic Dysfunction. A: Experimental design for B-cell blockade during CS exposure. *Clec2^KO^* mice were treated with anti-CD20 or isotype control antibody every 4 weeks starting at 4 weeks of age. Stating at 6 weeks of age, mice were exposed to daily CS exposure for 4 months prior to tissue harvest. B- G: H&E staining of lung tissue demonstrating TLOs after 4 month of CS exposure in *Clec2^KO^* mice treated with isotype control antibody (C,F), but not in the lungs of control (B,E) mice, or *Clec2^KO^* mice treated anti-CD20 antibody (D,G). H-J: H&E staining of lung tissue after 4 months of CS exposure in mice treated with anti-CD20 antibody (J) compared to isotype control (I), or control RA exposed mice (H). K: Western blot for soluble elastin in lung lysates from CS- exposed *Clec2^KO^* mice treated with isotype control or anti-CD20 antibody, as well as control mice. Blot for actin shown as a loading control. Scale bars = 100 µm.

### Increased Lymphatic Markers are Associated with Emphysema and Prominent TLOs among Patients with COPD

Our work demonstrated that lymphatic dysfunction in mice models many aspects of the autoimmune emphysema phenotype of COPD. Therefore, we next investigated whether a lymphatic signature could be identified in patients with COPD. We used spatial transcriptomics to analyze lung tissue from COPD patients with an emphysema phenotype and prominent TLOs compared to those with COPD without emphysema. We found increased expression of vascular markers including those associated with LECs (*Thy1*, *CD34*, *Col4a*) (31, 32) in the lung parenchyma of patients with severe emphysema compared to patients with COPD without emphysema (Figure 7a). Interestingly, we also found increased expression *NUDT4*, which has been recently identified as a marker of lymph node LECs (*33*) in the lungs of emphysema patients compared to those with COPD without emphysema. Furthermore, the lymphatic markers *Thy1* and *Ccl21* were associated with increased disease severity in patients with emphysema (Figure 7b, c). These results suggest that differences in lymphatic markers may be associated with a subset of COPD patients that are characterized by emphysema and TLOs.

**Figure 7:**
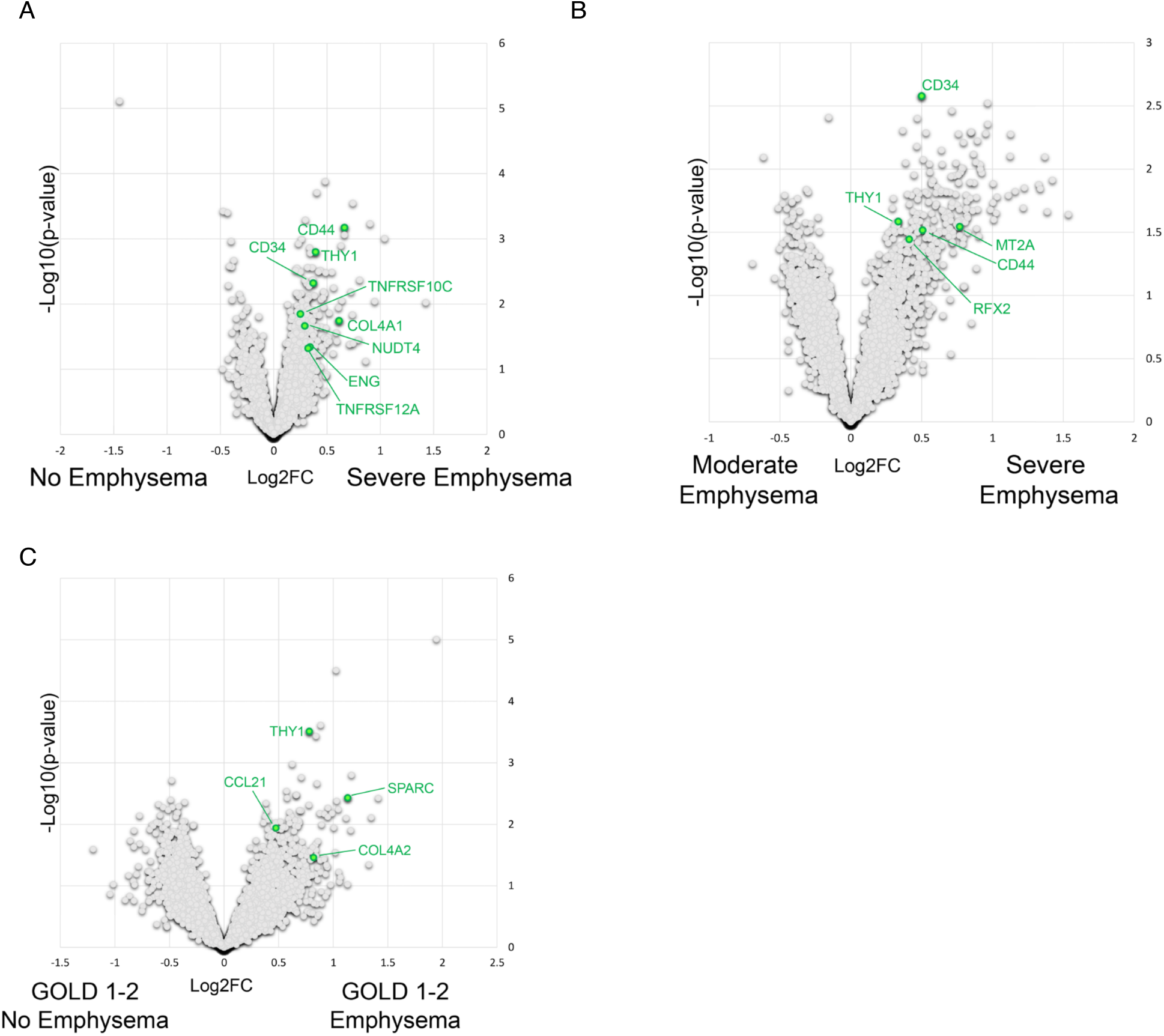
Emphysema phenotype and disease severity is associated with lymphatic markers in patients with COPD compared to patients without emphysema. A-C: FFPE lung sections from human COPD patients and controls were analyzed using Nanostring GeoMX Digital Spatial Profiler. Emphysema was assessed via CT-scan and classified according to the %LAA_950_. See Supplemental Table 1 for more information. A: Volcano plot comparing gene expression in the lung parenchyma of patients with severe emphysema (%LAA_950_ >17.7) compared to individuals without emphysema (%LAA_950_ <5.0). B: Volcano plot comparing gene expression in the lung parenchyma of patients with severe emphysema compared to moderate emphysema (%LAA_950_ 5-17.7) . C: Volcano plot comparing gene expression in the lung parenchyma of patients with GOLD 1-2 COPD patients with (%LAA_950_ >5.0) and without emphysema (%LAA_950_ <5.0).

## Discussion

COPD is a heterogeneous disease in which diverse endotypes reflect distinct pathologic mechanisms of disease. Among these, there is increasing recognition of a subset of patients with an emphysema phenotype that is characterized by a prominent B cell signature and lymphoid follicles (1, 2). Several studies have also suggested an autoimmune mechanism of disease in this subset of patients in which autoantibody production and self-reactive T cells are thought to drive lung injury (12–14, 34–36). While is clear that loss of tolerance and autoimmunity play a role in disease pathogenesis in a subset of COPD patients, the underlying factors that drive this mechanism of disease in some patients and not others has not been uncovered. In this study, we show that mice with lymphatic dysfunction develop autoantibodies in the lungs and increased emphysema after CS exposure. CS exposure leads to maturation of lung TLOs and increased autoantibody production in the lung tissue of these mice. Lymphatic dysfunction has been implicated in several autoimmune diseases including rheumatoid arthritis, systemic lupus erythematosus, and inflammatory bowel disease (19, 37). Our data support these previous studies and indicate that lymphatic dysfunction can also cause autoimmunity in the lung via the formation of autoantibody-producing TLOs. In addition, the data presented here support the intriguing hypothesis that underlying lymphatic dysfunction causes baseline self-reactivity that primes the lungs for further injury after CS exposure and drives an autoimmune endotype of COPD. This underlying lymphatic dysfunction may be due to genetic or developmental factors, or due to CS itself, which we have shown causes lymphatic dysfunction prior to the onset of lung injury in mice and is also seen in human emphysema (38).

Previous studies in both patients and animal models have shown lymphatic dysfunction and TLO formation in diverse settings of autoimmune and autoinflammatory disease (17, 19, 37, 39, 40). The work we describe here suggests that autoimmune emphysema may be just one of several manifestations of lymphatic impairment. Furthermore, our data showing that CS exposure results in TLO activation and increased autoantibody production may provide insights into autoimmune diseases with lymphatic dysfunction where CS exposure leads to worse disease, which has been observed in rheumatoid arthritis, Chron’s disease, and systemic lupus erythematosus (29, 41, 42). Whether lymphatic dysfunction results in a common mechanism of self-reactivity that is exacerbated by CS in several autoimmune diseases remains to be determined. Further studies are warranted to investigate whether CS exacerbates autoimmune disease by causing further lymphatic damage, by increasing maturation of self-reactive TLOs formed due to lymphatic dysfunction, or both.

We also have also used spatial transcriptomic analysis of human tissue to show increased expression of genes associated with the lymphatic vasculature in patients with an emphysema phenotype of COPD that is associated with TLOs. These patients are the subset most likely to have features of autoimmunity, and our data now provide the first evidence of differences in the lymphatic vasculature in the patients as well. Notably, it is unclear whether the increased presence of genes associated with lymphatics reflects a difference in lymphatic function in these patients, though it is interesting to note that we observe increased lymphatic vessel density in *Clec2^pltKO^* mice, likely reflecting the abnormal lymphatic morphology and architecture of these dysfunctional vessels (8). Interestingly, *Thy1*, which was found to be consistently associated both with emphysema phenotype and disease severity among patients with COPD, has been identified as a specific LEC marker in TLOs that specifies an LEC subset that promotes T cell function (43). Whether changes in lymphatic function is involved in the pathogenesis of disease in COPD patients with an autoimmune emphysema phenotype and prominent TLOs will be the subject of future investigations.

B cell blockade prevented TLO formation and decreased lung injury in mice with lymphatic dysfunction, providing strong evidence that lung injury in this model is due, at least in part, to the self-reactivity promoted by these structures. Though there are several lines of data suggesting that TLOs are pathogenic in COPD in both humans and animal studies (2, 4, 44, 45), defining their importance has been complicated by the fact that B cells and the adaptive immune system are dispensable for emphysema after CS exposure in mouse models (46). Indeed, a clinical trial of B-cell blockade in human COPD was terminated early due to increased risk of infection (5). However, among the clinically heterogeneous population of COPD patients, pathogenic TLOs are likely only a prominent marker of disease in a subset with an autoimmune emphysema phenotype. Recent data support this concept and have shed light on the clinical and histologic signatures of these patients (1, 2). Our data go further and for the first time suggest a connection between autoimmune emphysema and TLOs formed due to lymphatic dysfunction. Early identification of COPD patients with an autoimmune phenotype of emphysema as well as whether differences in lymphatic function may drive disease pathogenesis in these patients may lead to targeted therapies that are more likely to be effective.

The studies shown here take advantage of lymphatic dysfunction in *Clec2^pltKO^* mice that results from lack of separation between the blood and lymphatic systems and impaired lymphatic vessel maturation (9–11). While these mice also have defects in platelet count and function at baseline, the role of CLEC2 in hemostasis is minor and unlikely to contribute to the phenotype of TLO formation and lung injury that we observe (47–49). In addition, we and others have shown TLO formation associated with lymphatic dysfunction in other models (8, 18, 19), suggesting that it is lymphatic impairment that drives TLO formation in CLEC2-deficient mice. Interestingly, mice with a lymphatic leukocyte trafficking defect due to loss of CCR7, a cytokine receptor on leukocytes that binds CCL21 on the lymphatic endothelium, also develop lung TLOs (Supplemental Figure 2 and (15)). In addition, mice with loss of lymphatic FOXC2 develop lymphatic dysfunction that is also associated with TLOs (18). These data suggest that TLO formation in the lungs may be a common endpoint of lymphatic dysfunction caused by diverse anatomic and molecular mechanisms. Furthermore, single cell RNA sequencing analysis revealed that impaired lymphatic flow and lung TLOs in *Clec2^pltKO^* mice led to a subtype of LECs that is characterized by increased expression of MHC II antigen presentation genes. Interestingly, a similar LEC subtype was also identified in mice with lymphatic dysfunction due to loss of LEC FOXC2 that also develop TLOs (18). While typically seen in lymph nodes, these data suggest that this LEC subtype may arise outside of lymphoid tissue as a result of impaired lymphatic drainage from multiple causes. The mechanisms that lead to upregulation of antigen presentation genes in these LECs in setting of impaired flow are not entirely clear. Previous studies in other settings have shown that this LEC subset generally plays a tolerogenic role due to lack of co-stimulatory molecules (50, 51). Whether MHC II expression on lung LECs near TLOs affect the function of these structures during lung injury will be the subject of future investigations.

In summary, we have found that underlying lymphatic dysfunction in mice models an autoimmune emphysema phenotype of COPD. Though autoimmune emphysema has been the subject of previous investigations, to our knowledge this is the first study to support a direct role for lymphatic dysfunction in this setting. Identification of markers of lymphatic dysfunction in the subset of COPD patients with this phenotype may open the possibility of specific therapies targeted towards improving lymphatic function, inhibiting autoreactivity, or both.

## Methods

### Mice

*Clec2^flox^*, PF4Cre, and *Prox1-EGFP* (52) mice have been previously described(8, 11) and were maintained on a C57Bl/6 background. Mice were housed in the Weill Cornell animal facility in 12/12hrs light/dark cycles with ad libitum access to water and food. For all experiments, control and experimental animals were identically housed on the same rack in the animal facility. Both male and female mice were used in experimental and control groups

### Lymphatic Endothelial Cell Isolation for Flow Cytometry and Single Cell RNA Sequencing

To isolate lung LECs, mice carrying the *Prox1-EGFP* reporter (in which all LECs are labeled with GFP) were injected with AlexaFluor647-conjugated isolectin (Invitrogen) for intravital labeling of endothelial cells just prior to sacrifice, as previously described(53, 54). The lungs were then digested using dispase/collagenase in HBSS to generate a single cell suspension for FACS, with additional staining using antibodies for CD31 (Biolegend 102418), lineage cocktail (Bioloegend 133311), and Epcam (Biolegend 118225). LECs were sorted using positive selection for isolectin, CD31, GFP, and negative gating for Epcam and Lineage markers. For some studies, antibodies for mouse MHC II (Ebioscience 2450719) were also used for flow cytometry.

For single cell sequencing, (PROX1^+^, Pecam1^+^, Ptprc^-^, Lin^-^) LECs from *Clec2^pltKO^* (n=3) and control (n=3) mice were FACS isolated and sequenced on Illumina’s Next-generation sequencing platform. Sequencing reads in FASTQ format were aligned, and filtered feature-barcode matrices were generated for each sample using Cell Ranger (v.6.1.2). Using Seurat (55) cells were retained if they met the following criteria: less than 15% mitochondrial percentage, total counts below 10,000, and between 300 and 5,500 features. This approach was taken to exclude low-quality cells, lysed cells, and doublets. After normalization, 4,000 variable features were selected with the FindVariableFeatures function using the ‘vst’ method. FindNeighbors was then run after scaling using the first 20 principal components and a k.param of 15. Clustering was conducted using Seurat with a resolution of 0.2 and the default settings. Subsequently, UMAP reduction was applied utilizing the first 20 principal components.

Contaminating cell types (Ptprc^+^ or PROX1^-^, Cdh5^+^) were then removed and the remaining cells were renormalized and clustered based on the same parameters. Differential expression was performed using MAST (56) with random effects settings. The MHC II Protein Complex Binding (GO:0023026) Molecular Function gene signature, contributed by the Gene Ontology Consortium, was downloaded from Mouse MySigDb (56). After loading the analysis into Scanpy (57), functional enrichment scores for the signature were calculated for each cell using over representation analysis from decoupleR (58).

### Mouse Model of Cigarette Smoke Exposure and Emphysema

For cigarette smoke exposure studies, we used the inhalation exposure apparatus (TE- 10) by Teague Enterprises with 3R4F composition cigarettes (University of Kentucky Center for Tobacco Reference Products). Age matched mice beginning at 6–8 wk of age were exposed to CS (∼150 mg/m^3^) for a minimum of 3 hours per day, 5 days a week for 8 months(59). Age-matched mice exposed to room air were used as controls. RA control mice were identically housed in the same room, on the same rack, as CS exposed mice. To assess for emphysema, we calculated mean chord length (MCL). To ensure accurate representation of the tissue, we will use at least 8 randomly acquired 20x images from each mouse, using both the right and left lungs. We utilized morphometry software to quantify the length of chords within areas identified as airspace(60, 61). Using this method, it is possible to measure the size of the alveoli in all parts of the lung in a standardized and relatively automated manner(60, 61). Large airways, blood vessels, and other non-alveolar structures such as macrophages were manually removed from the images. The experimental group and genotype of the mice was blinded during acquisition and analysis of the images. For B-cell depletion, mice were injected with 250μg of purified rat anti-mouse CD20 (Utra- LEAF, Biolegend) or rat isotype control antibody retro-orbitally.

### Histology and Immunohistochemistry

Mice were sacrificed and tissue was perfused with PBS. Prior to harvest, lungs were inflated with 4% PFA at constant pressure of 25 cm H_2_O. Lungs were fixed in 4% PFA overnight at 4 degrees. The tissue was then dehydrated and embedded in paraffin for sectioning. 6-µm sections were H&E stained or immunostained with antibodies for: B220, CD3, CD138, and VEGFR3. All primary antibodies were incubated on slides overnight at 4 degrees. After washing, slides were incubated with AlexaFluor-conjugated secondary antibodies for 2 hours at room temperature. Slides were treated with DAPI-containing Vectashield and a cover slip was applied. Negative control slides were stained with secondary antibodies alone to control for autofluorescence of lung tissue. For detection of IgG deposition on mouse tissue, a AlexaFluor conjugated anti-mouse IgG antibody was used. Western blots were performed according to standard protocols and probed with anti-elastin (Abcam, ab21610) and anti-beta actin antibodies (Abcam, ab8226).

### Autoantibody and Cytokine Assays

Analysis of total IgA, IgG, and anti-collagen I and II in mouse BAL was done using a commercially available ELISA on a 96-well plate (Chondrex) and a plate reader. Anti-elastin antibodies were analyzed by ELISA made by coating a nickel-coated 96 well plates with His-tagged mouse elastin (MyBiosource, MBS2010590). Anti-elastin antibody from Abcam (217356) was used for standards. The plate was blocked with 1%BSA in PBS priory to incubation with mouse BAL or serial dilutions of anti-elastin standards for 1 hour. After washing, HRP-conjugated secondary antibodies were added to the plate and incubated for 1 hour. The plate was developed using TMB and read at 450/570nm. An autoantibody array to 128 antigens was performed on mouse BAL or lung lysates (UT Southwestern Genomics and Microarray Core Facility). The samples were treated with DNAse I, diluted and incubated with autoantigen array plate. The autoantibodies binding to the antigens on the array were detected with fluorescently labeled anti-IgG and anti-IgA antibodies and scanned with a GenePix® 4400A Microarray Scanner. The images were analyzed using GenePix 7.0 software to generate GPR files. The averaged net fluorescent intensity (NFI) of each autoantigen will be normalized to internal controls and was used to generate heat maps. Cytokine profiling of mouse lung homogenates was performed using a mouse proteomic cytokine profiler (R&D Systems) according to manufacturer’s instructions, which was performed in duplicate.

### Spatial Proteomics

Spatial Proteomics of murine TLOs was performed on FFPE murine lung sections with Nanostring GeoMX for Digital Spatial Profiling (DSP). First, adjacent H&E sections of lung tissue were used to identify TLO structures prior to Region Of Interest (ROI) selection. Second, slides were treated according to manufacturer’s instructions. Briefly, FFPE sections were deparaffinized, followed by antigen retrieval, and probe hybridization with photocleavable oligonucleotide tags. The proprietary proteomic panels were the following: Immune Cell Typing, Immune Activation Status, Myeloid, and Mouse Protein Core panels (Table 1). Once the slides were scanned, TLO’s were defined as a collection of >40 CD45+ cells (PMID: 36575294). Each ROI sample was collected and used to construct a library for sequencing using NextSeq500 Illumina platforms. Data processing was performed by the Weill Cornell Genomics Core. Proteomic data was audited, normalized and analyzed using R scripts for GeoMX DSP (‘GeomxTools’ (<u>10.18129/B9.bioc.GeomxTools</u>), ‘NanoStringNCTools’ (<u>10.18129/B9.bioc.NanoStringNCTools</u>), and ‘GeoMxWorkflows’ (<u>0.18129/B9.bioc.GeoMxWorkflows</u>) packages available from Bioconductor (https://www.bioconductor.org).

### Spatial Transcriptomics of Human Lung Tissue

For this study, we datamined the transcriptomic data obtained from 48 patients randomly selected for DSP transcriptomic profiling (Rojas-Quintero et al. AJRCCM 2023, in press). The cohort was comprised of 8 Non-Smoker controls (NSC), 13 ever-smoker controls, 17 COPD GOLD 1-2, and 10 COPD GOLD 3-4. Information regarding age, sex, smoking habit status (current vs. former), pack-years, and lung function (FEV1% and FEV/FCV) can be found in Table 2. FFPE lung sections from these subjects were treated according to manufacturer’s instructions. The Chest Imaging Platform software was used by two independent experts to determine the presence of emphysema from CT chest scans as the percentage ratio of low-attenuation areas below -950 Hounsfeld units in each lobe of lung (%LAA-_950_HU). Absence of emphysema was defined as LAA <5%. FFPE human lung sections were used for Nanostring GeoMX for Digital Spatial Profiling (DSP), according to manufacturer’s instructions. Briefly, FFPE sections were deparaffinized, followed by antigen retrieval, and probe hybridization with photocleavable oligonucleotide tags for whole transcriptome atlas (∼18,000 probes). Parenchyma ROI was defined as Pan-CK+ alveolar epithelial cells. RNA library was sequenced using NextSeq500 Illumina platforms. Data treatment was performed using R scripts for GeoMX DSP (‘GeomxTools’ (10.18129/B9.bioc.GeomxTools), ‘NanoStringNCTools’ (10.18129/B9.bioc.NanoStringNCTools), and ‘GeoMxWorkflows’ (0.18129/B9.bioc.GeoMxWorkflows) packages available from Bioconductor (https://www.bioconductor.org)). Significance was determined using mixed linear model set as log2 fold change cutoff +/-0.32 and p<0.05.

**Table 2.**
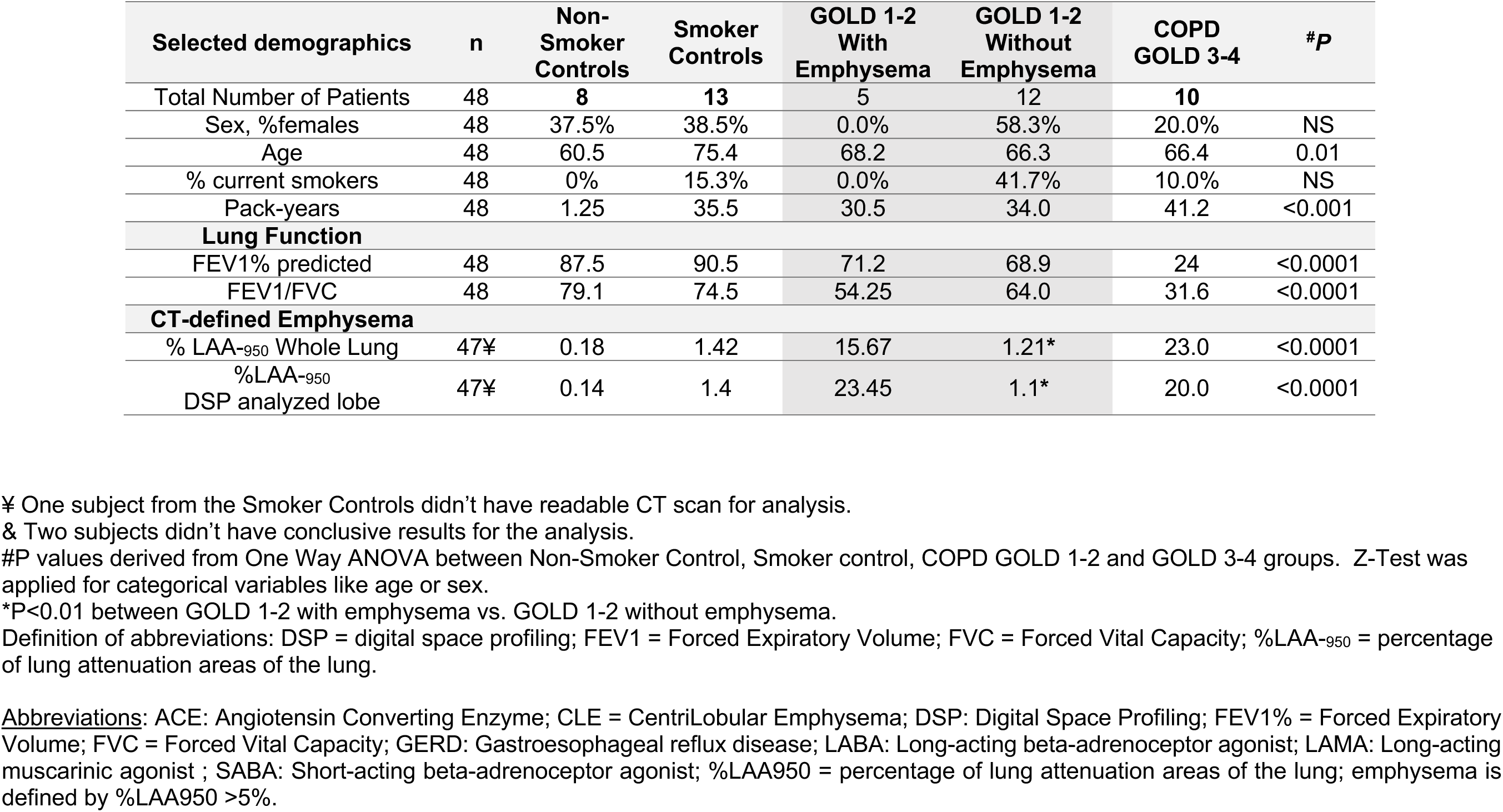
Selected demographics, lung function, and emphysema assessment.

### Statistics

Data are expressed as the mean ± SEM. Statistical significance was determined by unpaired, 2-tailed Student’s t-test or ANOVA using GraphPad Prism software. P values of less than 0.05 were considered statistically significant. Quantification of TLO number in lung tissue was performed using at least 5 randomly captured 10x images of H&E-stained lung sections per mouse representing the entirety of the lung tissue section for each sample. The genotype of the mice was blinded during quantification. TLOs were defined as discrete lymphocyte-dense accumulations on H&E-stained sections. Quantification of TLO area was performed using H&E sections of the entire lung of each mouse. TLOs were outlined by the free-drawing tool to calculate TLO area per lung section as a percentage of the total area of lung tissue in ImageJ. Quantification of lymphatic number in lung tissue was performed using at least 5 randomly captured 10x images of VEGFR3 staining per mouse.

### Study Approval

All animal experiments were approved by Weill Cornell Medicine Institutional Animal Care and Use Committee.

### Data Availability

RNA sequencing data have been deposited in GEO.

## Author contributions

B.S., K.K., A.T, T.L., and R.L. designed and conducted experiments, and analyzed data. T.L., S.H. and R.L. performed LEC isolation and single cell RNA sequencing analysis. T.P., J.R.Q., J.C-G., and F.P., conducted spatial proteomic analysis. J.R.Q. and F.P conducted spatial transcriptomic analysis. H.O.R. designed the research project, analyzed data, and drafted the manuscript.

## Acknowledgements

The authors thank Indu Raman and Chengsong Zhu at the UT Southwestern Microarray Core for assistance with autoantibody profiling. The authors also thank the Multiparametric In Situ Imaging (MISI) Core at WCM for technical assistance. This work was supported by K01HL145365 (H.R.), R01HL162990 (H.R.), the Robert Wood Johnson Foundation (H.R.), the Manning Foundation (H.R.), and the American Lung Association Innovation Award AAAALA2023 (H.R.).

## References

1. Sullivan JL, Bagevalu B, Glass C, Sholl L, KraA M, Martinez FD, et al. B Cell-Adaptive Immune Profile in Emphysema-Predominant Chronic Obstructive Pulmonary Disease. Am J Respir Crit Care Med. 2019;200(11):1434–9.

2. Polverino F, Seys LJ, Bracke KR, and Owen CA. B cells in chronic obstructive pulmonary disease: moving to center stage. Am J Physiol Lung Cell Mol Physiol. 2016;311(4):L687–L95.

3. Kheradmand F, Zhang Y, and Corry DB. Contributions of acquired immunity to the development of COPD in humans and animal models. Physiol Rev. 2022.

4. van der Strate BW, Postma DS, Brandsma CA, Melgert BN, Luinge MA, Geerlings M, et al. Cigarette smoke-induced emphysema: A role for the B cell? Am J Respir Crit Care Med. 2006;173(7):751–8.

5. Brusselle GG, Demoor T, Bracke KR, Brandsma CA, and Timens W. Lymphoid follicles in (very) severe COPD: beneficial or harmful? Eur Respir J. 2009;34(1):219–30.

6. Yadava K, Bollyky P, and Lawson MA. The formation and function of tertiary lymphoid follicles in chronic pulmonary inflammation. Immunology. 2016;149(3):262–9.

7. Aloisi F, and Pujol-Borrell R. Lymphoid neogenesis in chronic inflammatory diseases. Nat Rev Immunol. 2006;6(3):205–17.

8. Reed HO, Wang L, Sonett J, Chen M, Yang J, Li L, et al. Lymphatic impairment leads to pulmonary tertiary lymphoid organ formation and alveolar damage. J Clin Invest. 2019;129(6):2514–26.

9. Bertozzi CC, Schmaier AA, Mericko P, Hess PR, Zou Z, Chen M, et al. Platelets regulate lymphatic vascular development through CLEC-2-SLP-76 signaling. Blood. 2010;116(4):661–70.

10. Hess PR, Rawnsley DR, Jakus Z, Yang Y, Sweet DT, Fu J, et al. Platelets mediate lymphovenous hemostasis to maintain blood-lymphatic separation throughout life. J Clin Invest. 2014;124(1):273–84.

11. Sweet DT, Jimenez JM, Chang J, Hess PR, Mericko-Ishizuka P, Fu J, et al. Lymph flow regulates collecting lymphatic vessel maturation in vivo. J Clin Invest. 2015;125(8):2995–3007.

12. Nunez B, Sauleda J, Anto JM, Julia MR, Orozco M, Monso E, et al. Anti-tissue antibodies are related to lung function in chronic obstructive pulmonary disease. Am J Respir Crit Care Med. 2011;183(8):1025–31.

13. Wen L, Krauss-Etschmann S, Petersen F, and Yu X. Autoantibodies in Chronic Obstructive Pulmonary Disease. Front Immunol. 2018;9:66.

14. Byrne R, Todd I, Tighe PJ, and Fairclough LC. Autoantibodies in chronic obstructive pulmonary disease: A systematic review. Immunol Le@. 2019;214:8–15.

15. Fleige H, Bosnjak B, Permanyer M, Ristenpart J, Bubke A, Willenzon S, et al. Manifold Roles of CCR7 and Its Ligands in the Induction and Maintenance of Bronchus-Associated Lymphoid Tissue. Cell Rep. 2018;23(3):783–95.

16. Demoor T, Bracke KR, Vermaelen KY, Dupont L, Joos GF, and Brusselle GG. CCR7 modulates pulmonary and lymph node inflammatory responses in cigarette smoke- exposed mice. J Immunol. 2009;183(12):8186–94.

17. Winter S, Rehm A, Wichner K, Scheel T, Batra A, Siegmund B, et al. Manifestation of spontaneous and early autoimmune gastritis in CCR7-deficient mice. The American journal of pathology. 2011;179(2):754–65.

18. Gonzalez-Loyola A, Bovay E, Kim J, Lozano TW, Sabine A, Renevey F, et al. FOXC2 controls adult lymphatic endothelial specialization, function, and gut lymphatic barrier preventing multiorgan failure. Sci Adv. 2021;7(29).

19. Czepielewski RS, Erlich EC, Onufer EJ, Young S, Saunders BT, Han YH, et al. Ileitis- associated tertiary lymphoid organs arise at lymphatic valves and impede mesenteric lymph flow in response to tumor necrosis factor. Immunity. 2021;54(12):2795–811 e9.

20. Arroz-Madeira S, Bekkhus T, Ulvmar MH, and Petrova TV. Lessons of Vascular Specialization From Secondary Lymphoid Organ Lymphatic Endothelial Cells. Circ Res. 2023;132(9):1203–25.

21. Vandivier RW, and Ghosh M. Understanding the Relevance of the Mouse Cigarette Smoke Model of COPD: Peering through the Smoke. Am J Respir Cell Mol Biol. 2017;57(1):3–4.

22. Radder JE, Gregory AD, Leme AS, Cho MH, Chu Y, Kelly NJ, et al. Variable Susceptibility to Cigarette Smoke-Induced Emphysema in 34 Inbred Strains of Mice Implicates Abi3bp in Emphysema Susceptibility. Am J Respir Cell Mol Biol. 2017;57(3):367–75.

23. Litsiou E, Semitekolou M, Galani IE, Morianos I, Tsoutsa A, Kara P, et al. CXCL13 production in B cells via Toll-like receptor/lymphotoxin receptor signaling is involved in lymphoid neogenesis in chronic obstructive pulmonary disease. Am J Respir Crit Care Med. 2013;187(11):1194–202.

24. Bracke KR, Verhamme FM, Seys LJ, Bantsimba-Malanda C, Cunoosamy DM, Herbst R, et al. Role of CXCL13 in cigarette smoke-induced lymphoid follicle formation and chronic obstructive pulmonary disease. Am J Respir Crit Care Med. 2013;188(3):343–55.

25. Kelsen SG, Aksoy MO, Georgy M, Hershman R, Ji R, Li X, et al. Lymphoid follicle cells in chronic obstructive pulmonary disease overexpress the chemokine receptor CXCR3. Am J Respir Crit Care Med. 2009;179(9):799–805.

26. Di Stefano A, Caramori G, Gnemmi I, Contoli M, Bristot L, Capelli A, et al. Association of increased CCL5 and CXCL7 chemokine expression with neutrophil activation in severe stable COPD. Thorax. 2009;64(11):968–75.

27. Kaser A, Dunzendorfer S, Offner FA, Ludwiczek O, Enrich B, Koch RO, et al. B lymphocyte-derived IL-16 attracts dendritic cells and Th cells. J Immunol. 2000;165(5):2474–80.

28. Foy TM, Laman JD, Ledbetter JA, Aruffo A, Claassen E, and Noelle RJ. gp39-CD40 interactions are essential for germinal center formation and the development of B cell memory. J Exp Med. 1994;180(1):157–63.

29. Perricone C, Versini M, Ben-Ami D, Gertel S, Watad A, Segel MJ, et al. Smoke and autoimmunity: The fire behind the disease. Autoimmun Rev. 2016;15(4):354–74.

30. Packard TA, Li QZ, Cosgrove GP, Bowler RP, and Cambier JC. COPD is associated with production of autoantibodies to a broad spectrum of self-antigens, correlative with disease phenotype. Immunol Res. 2013;55(1-3):48–57.

31. Fiedler U, Christian S, Koidl S, Kerjaschki D, Emmett MS, Bates DO, et al. The sialomucin CD34 is a marker of lymphatic endothelial cells in human tumors. The American journal of pathology. 2006;168(3):1045–53.

32. Sauter B, Foedinger D, Sterniczky B, Wolff K, and Rappersberger K. Immunoelectron microscopic characterization of human dermal lymphatic microvascular endothelial cells. Differential expression of CD31, CD34, and type IV collagen with lymphatic endothelial cells vs blood capillary endothelial cells in normal human skin, lymphangioma, and hemangioma in situ. J Histochem Cytochem. 1998;46(2):165–76.

33. Takeda A, Hollmen M, Dermadi D, Pan J, Brulois KF, Kaukonen R, et al. Single-Cell Survey of Human Lymphatics Unveils Marked Endothelial Cell Heterogeneity and Mechanisms of Homing for Neutrophils. Immunity. 2019;51(3):561–72 e5.

34. Kheradmand F, Shan M, Xu C, and Corry DB. Autoimmunity in chronic obstructive pulmonary disease: clinical and experimental evidence. Expert Rev Clin Immunol. 2012;8(3):285–92.

35. Lee SH, Goswami S, Grudo A, Song LZ, Bandi V, Goodnight-White S, et al. Antielastin autoimmunity in tobacco smoking-induced emphysema. Nat Med. 2007;13(5):567–9.

36. Feghali-Bostwick CA, Gadgil AS, Otterbein LE, Pilewski JM, Stoner MW, Csizmadia E, et al. Autoantibodies in patients with chronic obstructive pulmonary disease. Am J Respir Crit Care Med. 2008;177(2):156–63.

37. Schwartz N, Chalasani MLS, Li TM, Feng Z, Shipman WD, and Lu TT. Lymphatic Function in Autoimmune Diseases. Front Immunol. 2019;10:519.

38. Summers B, Kim K, Clement CC, Khan Z, Thangaswamy S, McCright J, et al. Lung Lymphatic Thrombosis and Dysfunction Caused by Cigarette Smoke Exposure Precedes Emphysema in Mice. Scientific Reports. 2022;in press.

39. Rangel-Moreno J, Hartson L, Navarro C, Gaxiola M, Selman M, and Randall TD. Inducible bronchus-associated lymphoid tissue (iBALT) in patients with pulmonary complications of rheumatoid arthritis. J Clin Invest. 2006;116(12):3183–94.

40. Randolph GJ, Bala S, Rahier JF, Johnson MW, Wang PL, Nalbantoglu I, et al. Lymphoid Aggregates Remodel Lymphatic Collecting Vessels that Serve Mesenteric Lymph Nodes in Crohn Disease. The American journal of pathology. 2016;186(12):3066–73.

41. Harel-Meir M, Sherer Y, and Shoenfeld Y. Tobacco smoking and autoimmune rheumatic diseases. Nat Clin Pract Rheumatol. 2007;3(12):707–15.

42. Lakatos PL, Szamosi T, and Lakatos L. Smoking in inflammatory bowel diseases: good, bad or ugly? World J Gastroenterol. 2007;13(46):6134–9.

43. Shinoda K, Hirahara K, Iinuma T, Ichikawa T, Suzuki AS, Sugaya K, et al. Thy1+IL-7+ lymphatic endothelial cells in iBALT provide a survival niche for memory T-helper cells in allergic airway inflammation. Proc Natl Acad Sci U S A. 2016;113(20):E2842–51.

44. Roos AB, Sanden C, Mori M, Bjermer L, Stampfli MR, and Erjefalt JS. IL-17A Is Elevated in End-Stage Chronic Obstructive Pulmonary Disease and Contributes to Cigarette Smoke- induced Lymphoid Neogenesis. Am J Respir Crit Care Med. 2015;191(11):1232–41.

45. Seys LJ, Verhamme FM, Schinwald A, Hammad H, Cunoosamy DM, Bantsimba-Malanda C, et al. Role of B Cell-Activating Factor in Chronic Obstructive Pulmonary Disease. Am J Respir Crit Care Med. 2015;192(6):706–18.

46. D’Hulst A I, Maes T, Bracke KR, Demedts IK, Tournoy KG, Joos GF, et al. Cigarette smoke- induced pulmonary emphysema in scid-mice. Is the acquired immune system required? Respir Res. 2005;6:147.

47. Suzuki-Inoue K, Inoue O, Ding G, Nishimura S, Hokamura K, Eto K, et al. Essential in vivo roles of the C-type lectin receptor CLEC-2: embryonic/neonatal lethality of CLEC-2- deficient mice by blood/lymphatic misconnections and impaired thrombus formation of CLEC-2-deficient platelets. J Biol Chem. 2010;285(32):24494–507.

48. Hughes CE, Navarro-Nunez L, Finney BA, Mourao-Sa D, Pollitt AY, and Watson SP. CLEC-2 is not required for platelet aggregation at arteriolar shear. J Thromb Haemost. 2010;8(10):2328–32.

49. Rayes J, Watson SP, and Nieswandt B. Functional significance of the platelet immune receptors GPVI and CLEC-2. J Clin Invest. 2019;129(1):12–23.

50. GkounCdi AO, Garnier L, Dubrot J, Angelillo J, Harle G, Brighouse D, et al. MHC Class II Antigen Presentation by Lymphatic Endothelial Cells in Tumors Promotes Intratumoral Regulatory T cell-Suppressive functions. Cancer Immunol Res. 2021;9(7):748–64.

51. Santambrogio L, Berendam SJ, and Engelhard VH. The Antigen Processing and Presentation Machinery in Lymphatic Endothelial Cells. Front Immunol. 2019;10:1033.

52. Choi I, Chung HK, Ramu S, Lee HN, Kim KE, Lee S, et al. Visualization of lymphatic vessels by Prox1-promoter directed GFP reporter in a bacterial artificial chromosome-based transgenic mouse. Blood. 2011;117(1):362–5.

53. Barcia Duran JG, Lis R, Lu TM, and Rafii S. In vitro conversion of adult murine endothelial cells to hematopoietic stem cells. Nat Protoc. 2018;13(12):2758–80.

54. Nolan DJ, Ginsberg M, Israely E, Palikuqi B, Poulos MG, James D, et al. Molecular signatures of tissue-specific microvascular endothelial cell heterogeneity in organ maintenance and regeneration. Dev Cell. 2013;26(2):204–19.

55. Hao Y, Hao S, Andersen-Nissen E, Mauck WM, 3rd, Zheng S, Butler A, et al. Integrated analysis of multimodal single-cell data. Cell. 2021;184(13):3573–87 e29.

56. Finak G, McDavid A, Yajima M, Deng J, Gersuk V, Shalek AK, et al. MAST: a flexible statistical framework for assessing transcriptional changes and characterizing heterogeneity in single-cell RNA sequencing data. Genome Biol. 2015;16:278.

57. Wolf FA, Angerer P, and Theis FJ. SCANPY: large-scale single-cell gene expression data analysis. Genome Biol. 2018;19(1):15.

58. Badia IMP, Velez Santiago J, Braunger J, Geiss C, Dimitrov D, Muller-Dott S, et al. decoupleR: ensemble of computational methods to infer biological activities from omics data. Bioinform Adv. 2022;2(1):vbac016.

59. Cloonan SM, Glass K, Laucho-Contreras ME, Bhashyam AR, Cervo M, Pabon MA, et al. Mitochondrial iron chelation ameliorates cigarette smoke-induced bronchitis and emphysema in mice. Nat Med. 2016;22(2):163–74.

60. Laucho-Contreras ME, Taylor KL, Mahadeva R, Boukedes SS, and Owen CA. Automated measurement of pulmonary emphysema and small airway remodeling in cigarette smoke-exposed mice. J Vis Exp. 2015(95):52236.

61. Cloonan SM, and Choi AM. Mitochondria in lung disease. J Clin Invest. 2016;126(3):809–20.

