## Supplemental Data for "Lymphatic Dysfunction Models an Autoimmune Emphysema Phenotype of Chronic Obstructive Pulmonary Disease"

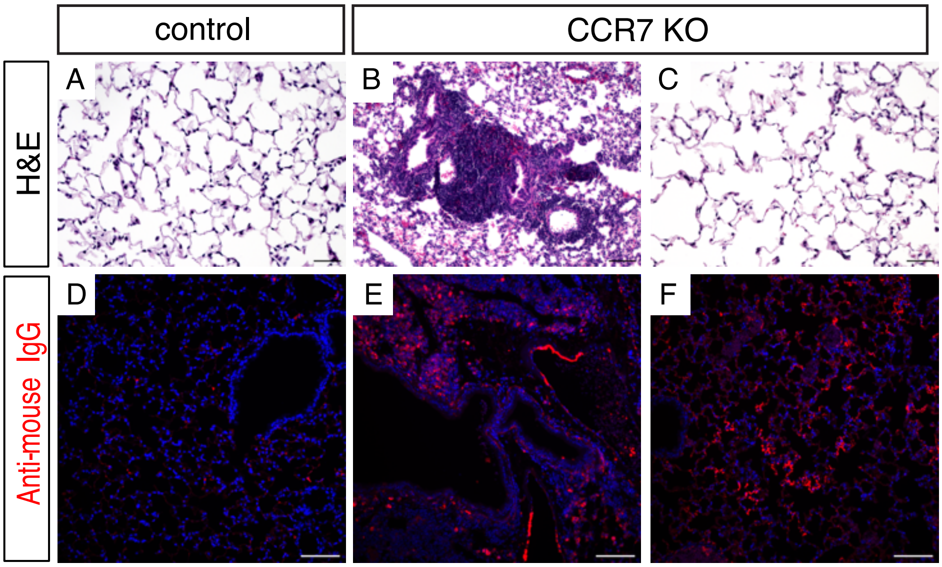


Supplemental Figure 1

A-C: H&E of lung sections from CCR7 KO (B,C) and control (A) mice. D-F: Staining of adult lung tissue from control (D) and CCR7 KO (E,F) for mouse IgG (red). Scale bars in A-C = 50 $\mu$m. Scale bars in D-F = 100 $\mu$m.

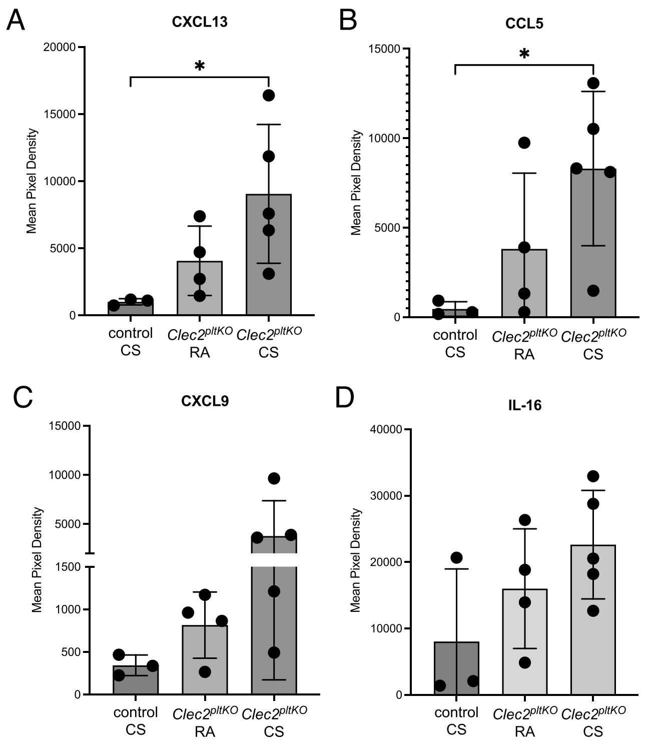


Supplemental Figure 2

A-D: Cytokine expression in whole lung lysates for CXCL13 (A), CCL5 (B), CXCL9 (C), and IL-16 (D) using a proteomic cytokine array performed in duplicate for all samples. All values are means ± SEM. *P* value calculated by ANOVA. $^*P < 0.05$.

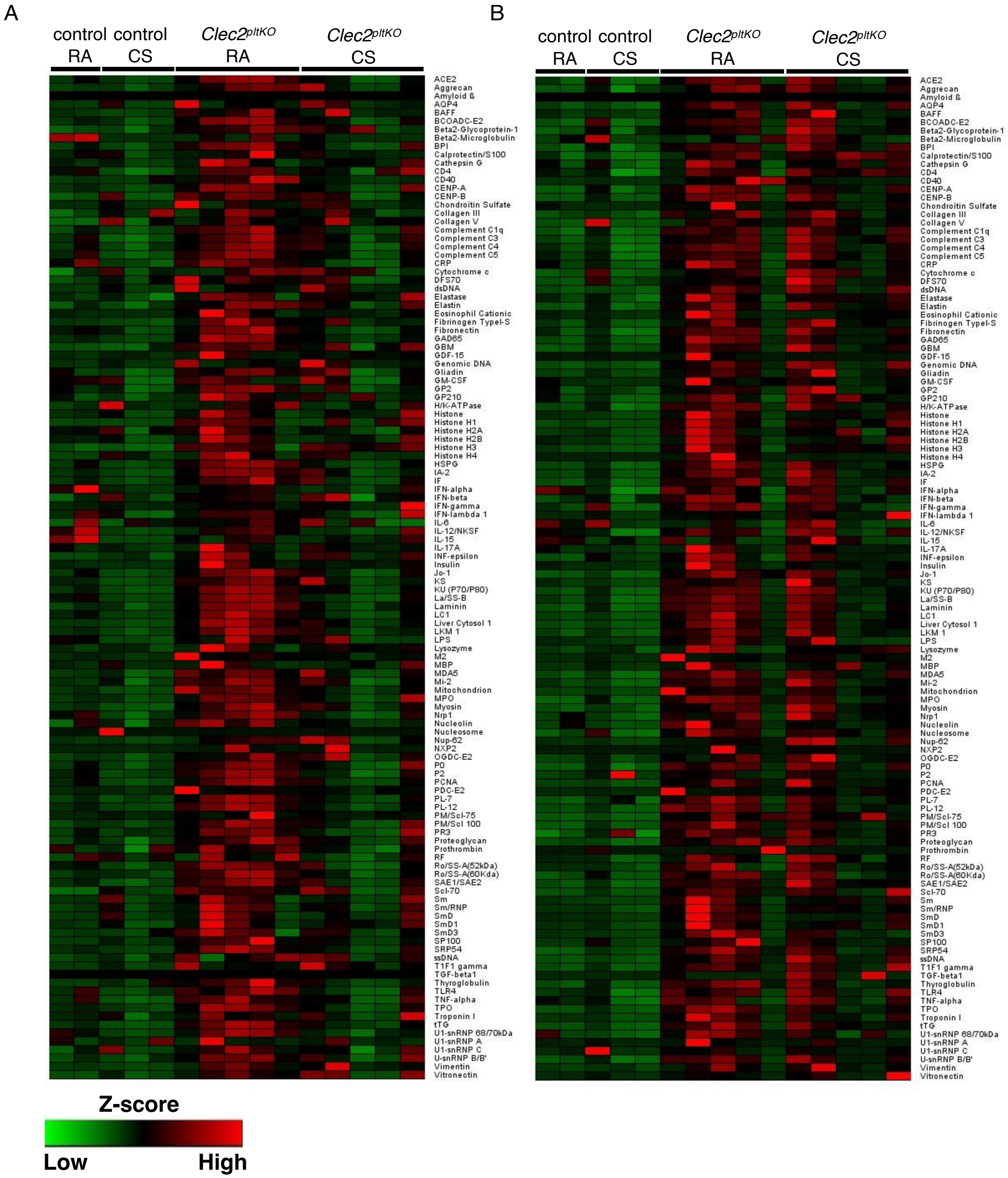


Supplemental Figure 3

A, B: Heatmap of IgG (A) and IgA (B) autoantibodies to 128 antigens detected by ELISA using BAL fluid from *Clec2*$^{pltKO}$ and control mice exposed to 8 months CS or RA and fluorescently-labelled mouse anti-IgG. Fluorescence intensity was normalized to internal controls.

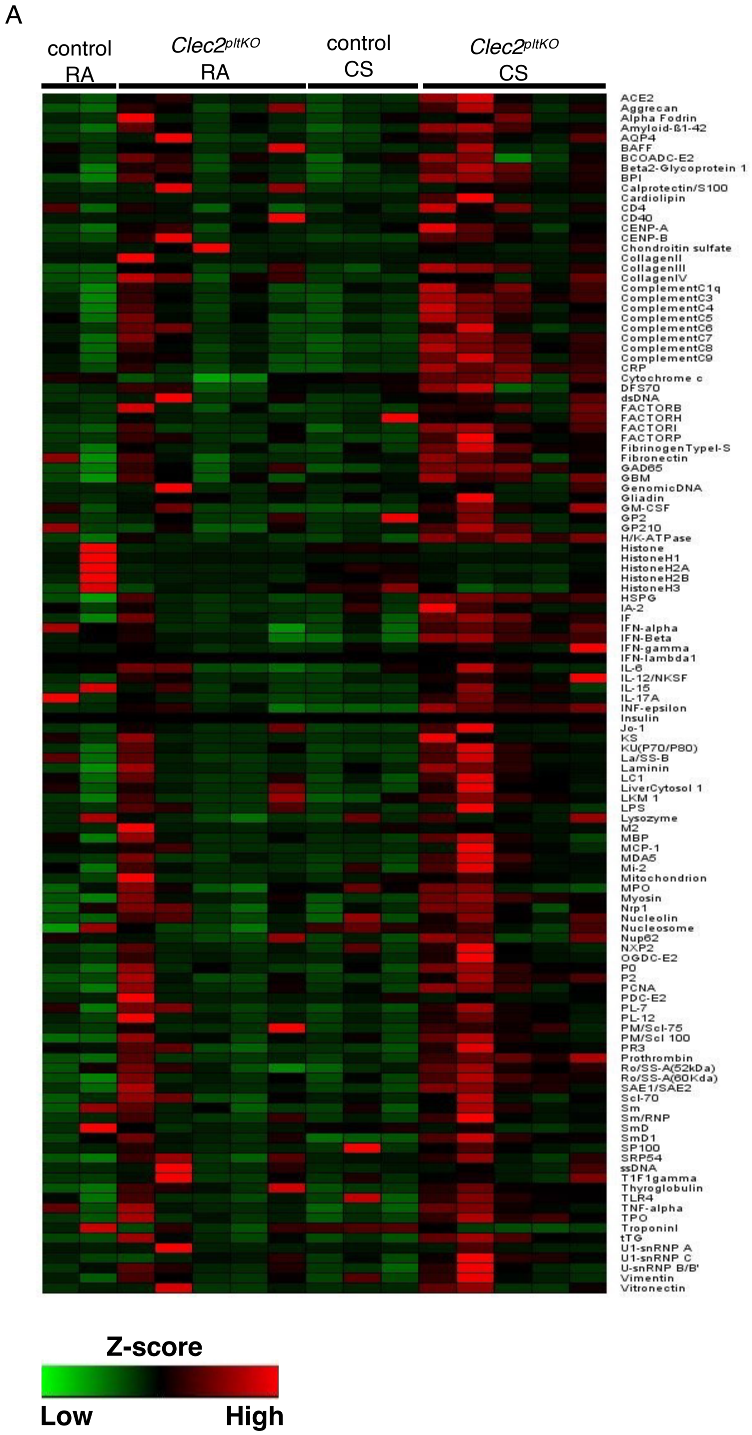


<u>Supplemental Figure 4</u>

A: Heatmap of IgA autoantibodies to 128 antigens detected by ELISA using lung lysates from *Clec2*$^{pltKO}$ and control mice exposed to 8 months CS or RA and fluorescently-labelled mouse anti-IgG. Fluorescence intensity was normalized to internal controls.

A

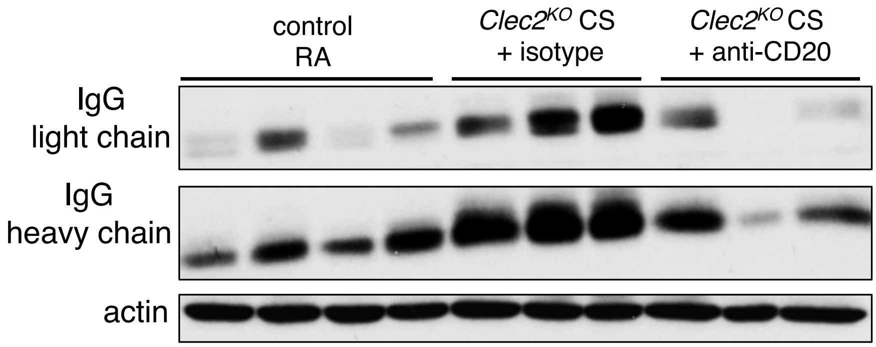


<u>Supplemental Figure 5</u>

A: Western blot for IgG antibody (light chain and heavy chain) in the lung lysate of mice exposed to 4 months of CS and treated with anti-CD20 or isotype control antibodies. Western blot for actin shown as a loading control.
